# Inferring Population Genetics Parameters of Evolving Viruses Using Time-series Data

**DOI:** 10.1101/437483

**Authors:** Tal Zinger, Pleuni S. Pennings, Adi Stern

**Affiliations:** Department of Molecular Microbiology and Biotechnology, School of Molecular Cell Biology and Biotechnology, Tel-Aviv University, Tel-Aviv, Israel; Department of Biology, San Francisco State University, San Francisco, California, USA

## Abstract

1

With the advent of deep sequencing techniques, it is now possible to track the evolution of viruses with ever-increasing detail. Here we present FITS (Flexible Inference from Time-Series) – a computational framework that allows inference of either the fitness of a mutation, the mutation rate or the population size from genomic time-series sequencing data. FITS was designed first and foremost for analysis of either short-term Evolve & Resequence (E&R) experiments, or for rapidly recombining populations of viruses. We thoroughly explore the performance of FITS on noisy simulated data, and highlight its ability to infer meaningful information even in those circumstances. In particular FITS is able to categorize a mutation as Advantageous, Neutral or Deleterious. We next apply FITS to empirical data from an E&R experiment on poliovirus where parameters were determined experimentally and demonstrate extremely high accuracy in inference. We highlight the ease of use of FITS for step-wise or iterative inference of mutation rates, population size, and fitness values for each mutation sequenced, when deep sequencing data is available at multiple time-points.

**Availability:** FITS is written in C++ and is available both with a highly user friendly graphical user interface but also as a command line program that allows parallel high throughput analyses. Source code, binaries (Windows and Mac) and complementary scripts, are available from GitHub at https://github.com/SternLabTAU/FITS.

## 2 Introduction

Evolutionary biology has traditionally relied on inferring evolutionary processes using data from one time point, namely from the present. With the advent of evermore accurate next generation sequencing (NGS) techniques, it is now possible to observe virus evolution in action – either through Evolve and Resequence (E&R) experiments (e.g., Acevedo et al., 2014; Foll et al., 2014; Stern et al., 2017) or from clinical samples obtained from patients (e.g., Dunn et al., 2015; Ramachandran et al., 2011; Zanini et al., 2015). Recent development of novel NGS techniques allow detection of ultra-rare alleles, even at frequencies of 10^−4^ or lower (Acevedo et al., 2014; Gelbart et al., 2018; Jabara et al., 2011; Lou et al., 2013; Meacham et al., 2011; Salk et al., 2018; Yang et al., 2013; Zhou et al., 2015). This allows tracking the fate of a mutation from the moment it is born, in particular RNA viruses that have high mutation rates.

The dynamics of allele frequency over time depends on the following factors: (a) the relative fitness of the allele as compared to the wild-type (WT) allele, denoted here as *w*, (b) the population-wide mutation rate *μ*, and (c) the population size (Figure 1A). These factors will lead to a *trajectory* of allele frequencies (Figure 1B). Low fitness of an allele is expected to manifest as a trajectory remaining at low frequencies based on mutation-selection balance, where for haploid populations we expect the frequency *f* to be equal to 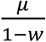. For example, a lethal mutation is ideally expected to be maintained at exactly the mutation rate *μ*. Population size will alter the extent to which random genetic drift affects the trajectory, with small populations being more susceptible to random fluctuations of frequencies than large populations. The mutation rate *μ* determines the rate at which an allele is introduced into the population at each generation.

**Figure 1.**
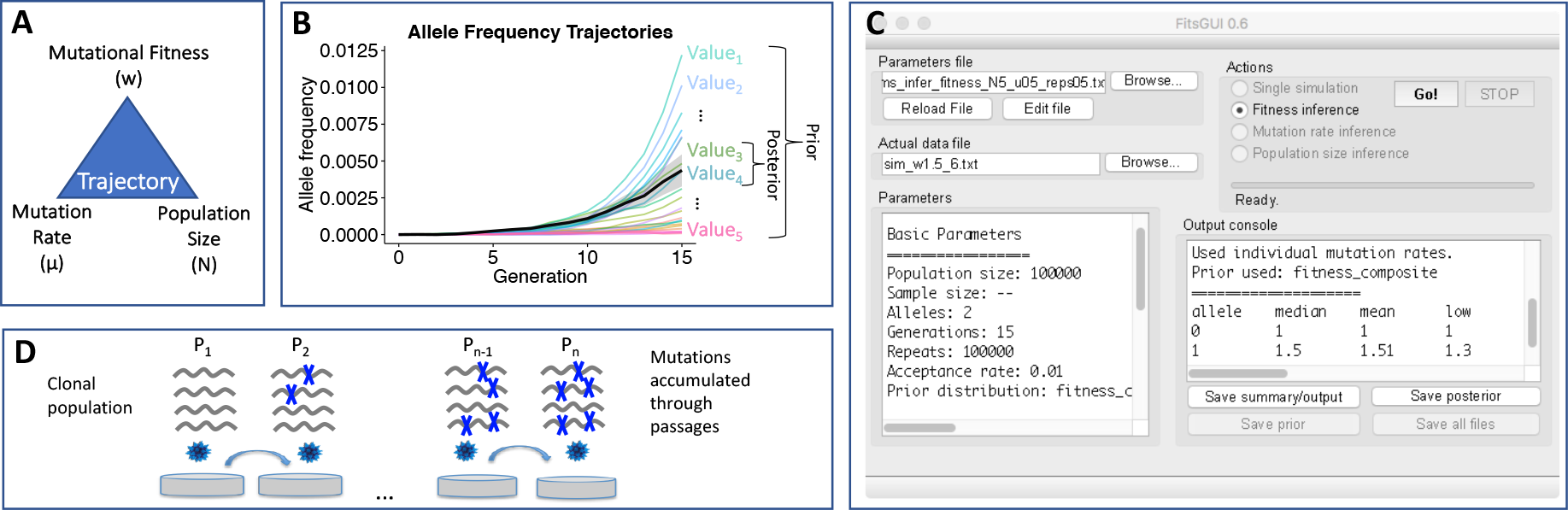
FITS overview (A) An allele frequency trajectory is determined by the mutational fitness, mutation rate, and population size. FITS can infer the value of one of these factors if information is present about the other two. (B) Rejection ABC in FITS works by repeatedly sampling values for the missing parameter from a prior distribution. These values are used to simulate the trajectories, which are then compared to the observed data (black line). The values used to generate the trajectories closest to the observed data (shaded area) constitute the posterior distribution, which ideally should be narrower than the prior. (C) FITS offers a user-friendly graphical user interface. The basic input required from the user is a file with allele frequency trajectories, and two of the three parameters: mutational fitness, population-wide mutation rate, and population size. Additional control of advanced parameters allows fine-tuning the results and inference. (D) FITS is designed to analyze time-series data, such as from evolve and re-sequence experiments. Here we show an experiment of serial passaging of viruses. A clonal population undergoes *n* passages, and mutations accumulate based on their fitness, based on the population-wide mutation rate, and based on the population size.

There have been many major advances in the development of approaches to infer selection and/or population size. Many such approaches chose to neglect the mutation rate, by assuming no recurrent mutation (Feder et al., 2014; Foll et al., 2014, 2015; Illingworth et al., 2012; Schraiber et al., 2016; Steinrücken et al., 2014), to ignore genetic drift (Acevedo et al., 2014; Renzette et al., 2014) or to allow only two alleles per locus (Bollback et al., 2008; e.g., Ferrer-Admetlla et al., 2016; Jónás et al., 2016; Terhorst et al., 2015; Topa et al., 2015). These simplifications may be problematic when studying the evolution of viruses, especially since diversity stems mainly from rapid mutation rates (Duffy et al., 2008), and since mutations emerge at low copy number and hence are dramatically affected by genetic drift. Moreover, recent advances in sequencing accuracy now allow tracking ultra-rare mutations from their birth over time (Acevedo et al., 2014), and have revealed that in virus populations four alleles often co-segregate at the same locus, most often at very low frequencies.

Given the trajectory of an allele, and given information on two of the three factors described above (fitness, population size mutation rate), it is possible to infer the third (Figure 1A). Here we introduce FITS, a highly user-friendly framework that allows inferring from time-series data either the fitness of a mutation, the mutation rate or the population size for a given virus population‥ FITS builds upon previous work (Foll et al., 2014), but incorporates several important additions such as assuming recurrent mutations, and allowing for the inference of mutation rates and population size. Particularly, the inference process may be done through a highly user-friendly graphical user interface (GUI, Figure 1C) that requires no expert previous knowledge. FITS is further available as a command-line tool, allowing parallel processing of genome-wide experimental data. Herein we have explored using both exact and noisy simulations the conditions where FITS yields informative inferences. We conclude that FITS is most often highly robust in classifying the fitness of an allele in conditions where input parameters are up to twice or half of the real values.

#### Box 1. Limitations of FITS

**FITS is designed to allow inference of fitness/mutation rate/population size from single-locus time-series allele frequency data. This is typical for next generation sequencing of virus populations, which is based on short reads, rendering the inference of haplotypes very difficult. FITS hence does not take into account linkage among sites. We hence recommend using FITS in the following conditions:**

- Viruses (or other microbes) with high recombination rates, where linkage is broken down rapidly. This is true for many (but not all) RNA viruses (Worobey and Holmes, 1999).
- E&R experiments that are based on a limited number of generations. For example, given a mutation rate of 10^−5^ for poliovirus and a clonal population at the beginning of the experiments, we expect that >90% of the virus genomes will bear only one mutation after seven replication cycles, and >70% after fourteen (as explored herein). Notably, for organisms with lower mutation rates this will be true for many more generations

**We recommend avoiding the use of FITS if:**

- There is a high probability that there are many linked mutations on the same genome
- The number of generations between the sampled time points is unknown
- The data is unreliable (for example, low coverage and hence low copy number of a mutation)
- The data is available for no more than two time points

## 3 Inferring Parameters Using ABC

### 3.1 Overview

Our method relies on the rejection ABC method, which gained popularity in recent years (Beaumont, 2010; Csilléry et al., 2010; Sunnåker et al., 2013). We start off with empirical data of allele frequencies over time, which can be derived, for example, from sequencing of a virus population during a serial passaging experiment (Figure 1D). Data from these experiments may or may not be from consecutive generations. Possible values for the parameter in question (fitness, mutation rate, or population size) are sampled from an appropriate *prior distribution*, and used for simulating *trajectories* (Figure 1B), starting at the frequencies observed at the first time-point. We then use the *ℓ*_1_ distance as a distance function between the trajectories, while using allele frequencies in generations shared by the observed and simulated trajectories as summary statistics. Namely, for each generation *g* ∈ {1 … *n*} that is shared between the simulated and observed data, the *sim*ulated frequency of each allele *a* ∈ {1 … *k*} is subtracted from the *obs*erved frequency. The individual distances are then summed to a scalar value representing the distance: 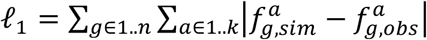. The top 1% of simulations most similar to the empirical data (with the lowest *ℓ*_1_ distance) are used as an approximation of the *posterior distribution* relevant to the inferred factor (Foll et al., 2014; Sunnåker et al., 2013). From this distribution, we take the median as a point estimate for the inferred factor (Aeschbacher et al., 2013; Csilléry et al., 2010; van der Vaart et al., 2015).

Simulated trajectories are generated using the two-step Wright-Fisher with selection model. The two steps of the simulation account for the fitness of an allele and for random genetic drift. The first step may be thought of as corresponding to a viral population multiplying in the cell, with viruses with preferred alleles (higher fitness) creating more progeny. That is, the proportion of the mutant allele at time *t* is first calculated as *P*_*t*_ = *w* ⋅ *x*_*t*−1_ ⋅ *μ*_*b*_ + 1 ⋅ (1 − *x*_*t*−1_) ⋅ *μ*_*f*_, where *w* is the fitness of the mutant allele, *x*_*t*_ is the mutant’s frequency at time *t*, and *μ*_*b*_ and *μ*_*f*_ are the backward and forward mutation rates, respectively. Next, random genetic drift is applied. For viruses, this may be thought of as the process through which progeny virions “hit” their target cell by chance, regardless of their genetic content. The proportion of the mutant allele at time *t* in the upcoming generation is thus calculated by applying binomial sampling: 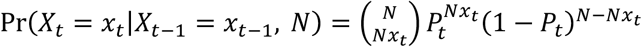, where *N* is the population size. Similar analogies can be made for other organisms.

When performing serial passaging, a population bottleneck may be imposed upon the population every generation or every few generations (Figure 1D), based on the user input. Furthermore, only a fraction of the genomes may be sampled for sequencing. FITS is able to account for both these bottlenecks by performing additional binomial sampling steps. We note that all in all, genetic drift may have a critical effect on the trajectory of an allele. This will be manifested in two ways: first via “classical” genetic drift that occurs when the population size is low. Stochastic effects will also exert their effect when the copy number of a new mutation is very low. This may occur in real-life infections that begin with a single virion (Keele et al., 2008) or in an experimental evolution setup beginning with a clone (Figure 1D). In both cases, a novel mutation will enter the population at an initially very low frequency and at a very low copy number.

In order to verify that the posterior distribution indeed gives a meaningful estimate we test whether the ABC process yields a posterior distribution that is significantly narrower than the prior distribution (see Figure 1B). For this purpose we apply Levene’s test on the prior and posterior distributions (van der Vaart et al., 2015). In the next section, we will discuss how FITS can be used to infer (1) fitness, (2) mutation rates, or (3) population size.

### 3.2 Inferring fitness values

The relative mutational fitness (*w*) of an allele is a measure of its *advantageous* (*w* > 1), *deleterious* (0 ≤ *w* < 1) or *neutral* (*w* = 1) effect on the reproductive success of the allele-bearing individuals, relative to individuals with the wild-type (WT) allele (for which *w* = 1). For our purpose the interval of *w* ∈ [0,2] is used. FITS may assume a uniform prior distribution in this interval, but as fitness tends to be biased towards deleterious mutations (Huber et al., 2017; Sanjuán, 2010), we allow a composite prior (see Figure S1) which is aimed at increased sampling of the deleterious range, to provide adequate representation in the process. This recapitulates the importance of choosing an appropriate prior in order to infer values more accurately (Sunnåker et al., 2013). Finally, as described above, the median of the posterior distribution is outputted as an estimate of the allele’s fitness.

### 3.3 Inferring fitness category

Often a user may be less interested in the exact fitness value, but will be more interested in broadly classifying an allele as deleterious, advantageous, or neutral. Therefore, we define an allele as *advantageous* (ADV) if at least 95% of its posterior distribution is greater than one, *deleterious* (DEL) if at least 95% of its posterior distribution is smaller than one and *neutral* (NEU) if at least 95% of its posterior distribution is equal to one. This classification is based on the measure of significance for a posterior distribution previously proposed (Beaumont and Balding, 2004), referred to as a “Bayesian p-value” (Foll et al., 2014). We note that classifying an allele as neutral is technically challenging, since the posterior distribution likely includes values near one but not equal to one. When more than 50% (but less than 95%) of the posterior distribution is positioned within the appropriate interval, FITS ambivalently classifies the allele as ?ADV, ?NEU, or ?DEL.

### 3.4 Inferring Mutation Rates

Many models utilize a single *mutation rate μ* to describe the probability of an allele to change, i.e. for every single allele *μ* = Pr(*A*_*1*_ → *A*_*j*≠*i*_). Nevertheless, several studies have recently shown that mutation rates may vary between different pairs of alleles (Abram et al., 2010; Acevedo et al., 2014; Zanini et al., 2017). FITS has been therefore designed to infer the mutation rate between any pair of alleles separately, given input on the fitness of the allele and the population size. FITS samples a value for the exponent (*n*) from a uniform prior, such that 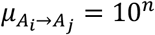. Setting the range of the prior to [−7,−2] captures most mutation rates of viruses (Acevedo et al., 2014; Sanjuán et al., 2010). This range can be reconfigured when applied to other organisms.

### 3.5 Inferring Population Size

Population size (*N*) affects the extent to which genetic drift exerts an effect on the allele frequency trajectory, and hence the degree to which fluctuations occur in the frequency of the allele. In the absence of genetic drift, the allele trajectory is expected to be monotonic. Once again, we sample the exponent (*m*) from a uniform distribution, such that *N* = 10^m^. A range of [3,8] should capture many experimental evolution settings, but this range can be configured by the user. In the case of population size, the aim of FITS is not necessarily to give an accurate estimate of *N*, but rather to give bounds on this parameter. This is because, for example, for large values of *N* the allele frequency trajectory will be very similar regardless of the precise value of *N*. In such a case, FITS will report that the posterior distribution spans a certain interval of values, the lowest of which can be interpreted as a lower bound on *N*. We emphasize here that the use of more than one locus to infer the mutation rate or the population size will obviously lead to a more robust inference. Currently FITS supports analysis on single loci only. By analyzing one by one the set of all known neutral alleles, we suggest that a distribution of inferred mutation rates or population sizes may be obtained, yielding more information than analysis of one locus only.

### 3.6 Graphical User Interface

In order to make FITS accessible to the wide scientific community, we developed a GUI that allows user to infer fitness/mutation rates/population size with a click of the button (Figure 1C). This GUI is cross-platform, and written in C++, using the Qt framework (https://www.qt.io). Binaries are available for Windows and Mac, but may be compiled on other platforms as well. The basic setup requires giving as input two of the three parameters in the *parameters file*, inputting the observed allele frequency trajectory in the *actual data file*, and pressing “*Go!*” (see Figure 1C).

## 4 Accuracy of FITS

### 4.1 Theoretical accuracy

As mentioned above, we use Levene’s test to compare the prior and posterior distributions, to show whether the point estimate given by FITS is meaningful. We generated datasets of simulated biallelic frequency trajectories over 15 generations, with different combinations of parameter values: *N* = {10^4^, 10^5^, 10^6^}, *μ* = {10^−6^, 10^−5^, 10^−4^} and *w* = {0, 1.0, 1.5}. Simulations all began from an initial frequency of zero, mimicking a situation when a new mutation is born in an initially clonal population. For each such combination, 100 replicates were generated, yielding a total of 2,700 datasets. We then used FITS to infer the fitness of the mutant allele from these datasets, and tested the reliability and precision of FITS.

We began by analyzing how many datasets yielded a posterior distribution significantly narrower than the prior distribution based on Levene’s test, indicating that FITS gained information from this dataset. Our results show that inference tends to be most reliable when *Nμ* ≥ 1 with 99-100% of datasets analyzed yielding a narrowed posterior (Figure 2A). On the other hand, when *Nμ* < 1, a much smaller portion of datasets were narrowed. This most likely derives from the fact that when *Nμ* < 1, a new mutation may not be created at all, and allele frequencies will remain at zero for most generations, regardless of the fitness. Therefore, we introduced a warning into FITS that states that for *Nμ* < 1 inference may not be reliable. We next turned to testing the effect of the number of simulations (i.e., the number of samples from the prior distribution) on reliability of inference (Nakagome et al., 2013). Indeed, we saw significantly more narrowed posteriors when increasing the number of simulations above 10^4^ (t-test, p≪0.01). However, increasing from 10^5^ to 10^6^ simulations did not lead to a significant improvement (t-test, p=0.27, Figure 2B).

**Figure 2.**
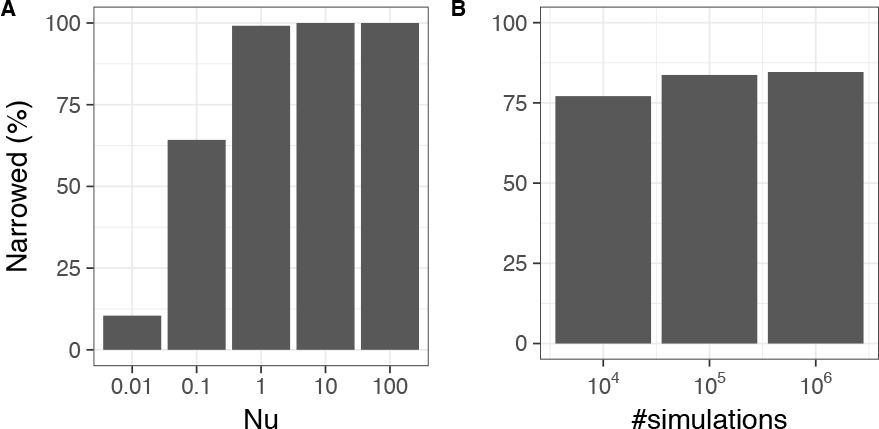
Analysis of simulated data (parameter values described in the text) and the ability of FITS to infer meaningful information: (A) The percent of datasets where the posterior distribution is significantly narrower than the prior distribution (Levene’s test with p<0.05) is shown to depend on the product *Nμ*. (B) The number of samples from the prior distribution (number of simulations) is shown to have a small effect in improving the percent of datasets with a narrower posterior distribution.

### 4.2 Accuracy of fitness estimates

After establishing that reliable results are obtained by the combination of (a) *Nμ* ≥ 1, corresponding to a large enough copy number of the allele, and (b) 10^5^ simulations per dataset, we went on to test the accuracy of point estimates given by FITS using the biallelic model. A subset of the simulated data described above was chosen such that *N* = 10^5^ and *μ* = 10^−5^, and the posterior distribution was narrower than the prior distribution (Levene’s test, p<0.05). Results of FITS were found to be quite satisfactory: the fitness of lethal alleles (true fitness equals to 0) was inferred as 0.07±0.11 (mean and standard deviation), the fitness of neutral alleles (true fitness equal to 1) was inferred as 0.91±0.16 and the fitness of advantageous alleles (true fitness equal to 1.5) was inferred as 1.44±0.12 (Figure 3A). Thus, the fitness of lethal alleles was slightly overestimated whereas neutral and advantageous alleles were slightly underestimated.

**Figure 3.**
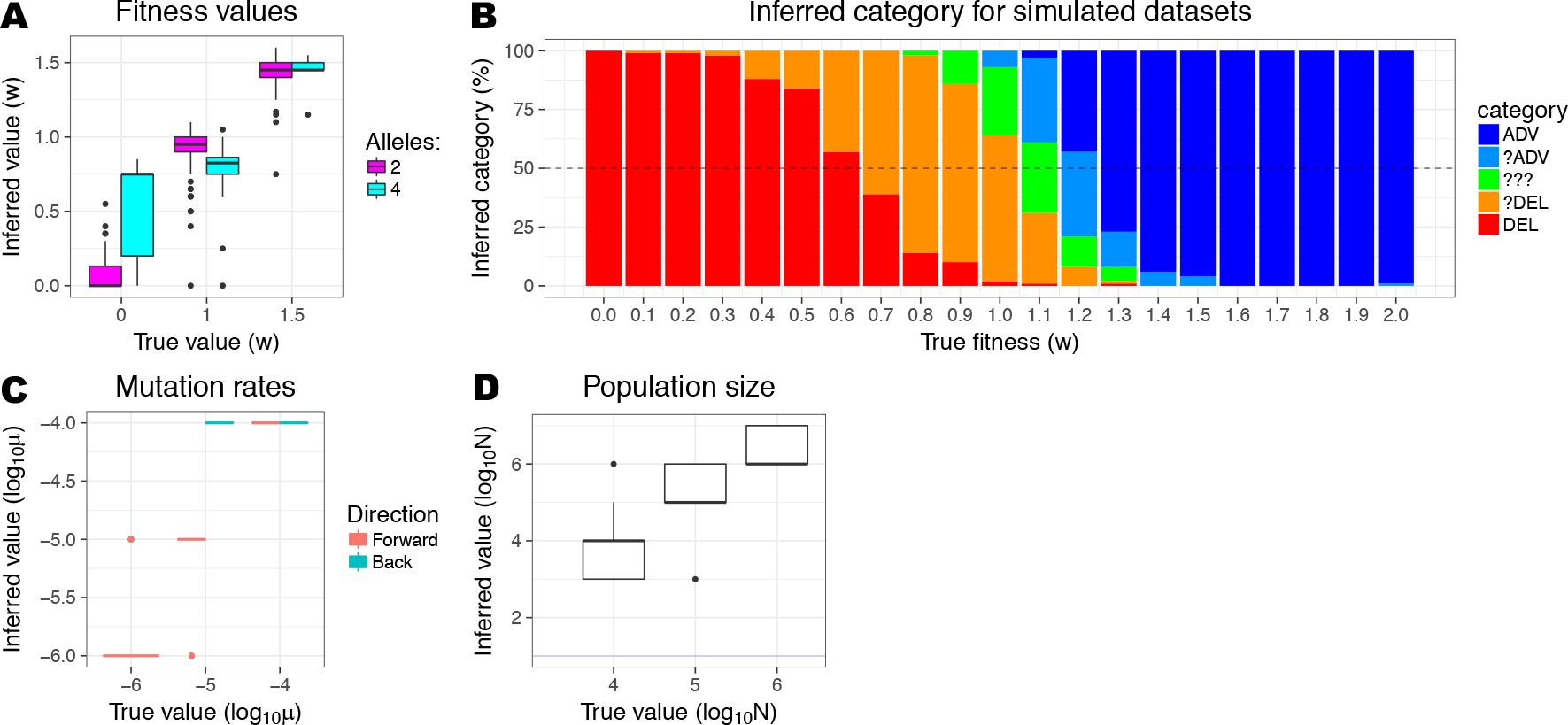
Results of FITS on simulated data. (A) Accuracy of fitness inference for simulated biallelic and quadrallelic datasets. (B) Accuracy of FITS in classifying alleles on 100 simulated biallelic datasets. For each fitness value, we show how many datasets were classified in each category. (C) Accuracy of back and forward mutation rate inference. No posterior distributions were narrowed for back mutation of *μ* = 10^−6^ and hence estimates are not shown. (D) Accuracy of population size inference. Boxplots in (A), (C), and (D) display boxes spanning the lower and upper quartiles (interquartile range, IQR). The middle band represents the median, and whiskers extend to 1.5 times the IQR. Points beyond this range are outliers.

We next sought to test the performance of FITS using the quadrallelic model (using four alleles), selecting biologically-plausible sets of fitness values (Table S1), with different mutation rates for transitions (10^−5^) and transversions (10^−6^), based on viruses where the transition/transversion ratio is often around ten (Illingworth et al., 2014). In 82 of 600 cases, the posterior distributions were narrowed for all alleles, and further analyzed. In general, FITS inferred the fitness of lethal alleles as 0.56±0.32, the fitness of neutral alleles as 0.79±0.18 and the fitness of advantageous alleles as 1.42±0.16. The accuracy of inference was strongly affected by the nature of an allele, with fitness of transition alleles inferred more accurately (Figure S2). This could be once again attributed to low copy number of the transversion alleles.

Our results show a general tendency to overestimate the fitness of lethal alleles (*w* = 0). This is most likely since zero represents the edge of the fitness interval (we cannot estimate values lower than zero), and our point estimate is the median of the posterior. On the other hand, fitness values of neutral and advantageous alleles are slightly underestimated. This effect seems to be related to the stochastic effects of copy number: in the initial generations, the copy number of a newly born mutation is almost always very low (depending on *N*), and thus an allele may be lost and regenerated over several generations till it “takes off” due to selection. This will lead to lower allele frequencies in general, which will resemble simulations with lower fitness values.

### 4.3 Classifying allele fitness

Although FITS gives a point estimate as an output, for many researchers, the category of the allele’s fitness (DEL, NEU, ADV) is more important than the exact value. We thus set out to see how accurately FITS categorizes alleles. In order to do so, we generated datasets by simulating trajectories for different fitness values with fixed population size (*N* = 10^5^) and mutation rate (*μ* = 10^−5^). For each fitness value, we generated 100 replicates and used FITS to infer the fitness of the mutant allele (Figure 3B). Our results showed that in general, FITS is able to quite accurately classify the allele, in particular when including the ambivalent ?ADV, ?NEU, ?DEL labels as well. In addition, we found that the mutant allele was classified as advantageous only when it was indeed so. Only a few datasets (7/100) yielded the ?ADV ambivalent labeling, and none yielded ADV when the actual fitness value was *w* ≤ 1. This is in consistent with FITS’ conservative estimation of advantageous alleles, which is a desired behavior for many users.

### 4.4 Mutation rate accuracy

In order to infer the mutation rate, one must begin by studying an allele whose fitness is known. This may be assumed to be the case for synonymous mutations, which are most often neutral, or for an allele where external information is available regarding fitness. Here we tested the ability of FITS to infer mutation rates given a neutral allele, by simulating datasets with varying mutation rates, while retaining *N* = 10^5^ and *w* = 1.0. Forward and back mutation rates were set to be the same value. Not surprisingly, FITS was mostly unable to infer the back mutation rate, as manifested in only 64 of 300 narrowed posterior distributions for the back mutation (compared to 300/300 for the forward mutation), and inexact point estimates (Figure 3C). This is likely because in our context, back mutations operate on the mutant allele, which exists in a very low copy number. Mean values for the forward (back) mutation rate measured −4±0 (−4±0), −5.07±0.26 (−4±0) and −5.92±0.27 for log_10_ *μ* = −4, −5, −6 respectively. These results recapitulate the findings above that show that when *Nμ* or the allele copy number are low (as occurs here when the mutation rate is 10^−6^), inference is less reliable.

### 4.5 Population size accuracy

The fitness of the allele (as well as the mutation rate) must be also known in order to infer the population size. Once again, we here mimicked inference given a neutral allele, by simulating *w* = 1 and *μ* = 10^−5^ over 100 datasets, and inferred the population size using FITS (Figure 3D). In terms of narrowed posterior distributions, we got fractions of 100/100, 98/100 and 87/100 for *N* = 10^4^, 10^5^, 10^6^ respectively. Our point estimates of population size were: 3.71±0.66, 5.41±0.55 and 6.49±0.5 for log_10_ *N* = 4,5,6 respectively. The wider variance obtained here shows that FITS will be useful in putting bounds on the population size but may be less useful in inferring the precise population size value.

### 4.6 Example for iterative approach

Given genomic sequencing data, FITS can be used iteratively to determine population size, mutation rate and fitness values for all alleles. We suggest the workflow illustrated in Figure 4.

**Figure 4.**
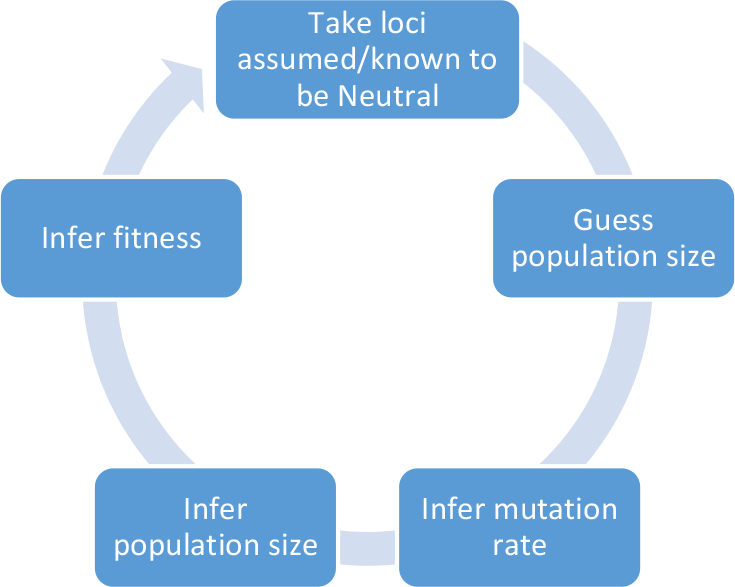
Iterative approach to determine parameters of a population.

As an example, we take 100 simulated biallelic loci, in which the mutant allele is neutral (*w* = 1). This mimics a partial set of synonymous mutations in a viral genome. We guessed different population sizes (10^4^, 10^5^, 10^6^), and inferred the mutation rate for each locus. Ultimately, we inferred a genomic-wide mutation rate as the median of inferred mutation rates from all loci (Table 1). FITS consistently (and correctly) inferred 10^−5^ as the mutation rate regardless of the guessed population size.

**Table 1.**
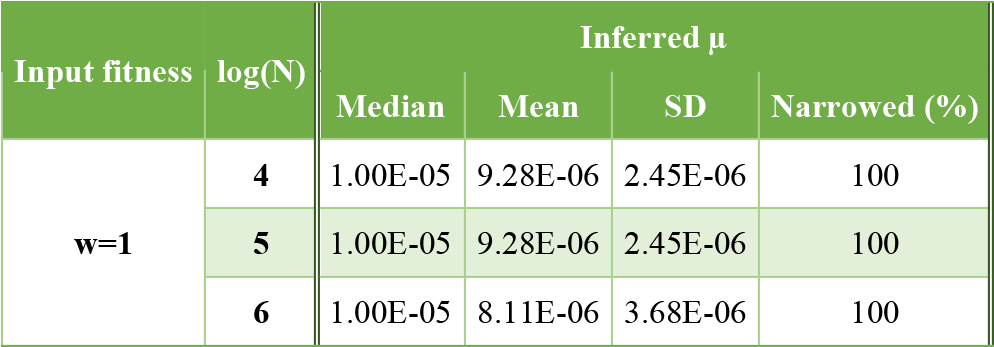
First iteration – assuming several population sizes and allele neutrality (w=1.0).

| Input fitness | log(N) | Inferred $\mu$ | | | |
| --- | --- | --- | --- | --- | --- |
|  |  | Median | Mean | SD | Narrowed (%) |
| $w=1$ | 4 | 1.00E-05 | 9.28E-06 | 2.45E-06 | 100 |
|  | 5 | 1.00E-05 | 9.28E-06 | 2.45E-06 | 100 |
|  | 6 | 1.00E-05 | 8.11E-06 | 3.68E-06 | 100 |

Given the inferred mutation rate of 10^−5^, we set out to infer the population size (Table 2**Error! Reference source not found.**). Two out of 100 datasets gave insignificant results. The median of the inferred population size across all datasets was 10^5^, again in line with the real value.

**Table 2.** First iteration – inferring population size given the inferred mutation rate

| Input $\mu$ | Median N | Mean N | SD N | Narrowed (%) |
| --- | --- | --- | --- | --- |
| $10^{-5}$ | 1.00E+05 | 4.94E+05 | 4.50E+05 | 98 |

Given the above population size and mutation rate, the fitness of the mutant allele was inferred, yielding a range of fitness values, most of which were very close to 1.0 (Figure 5). As we show below, this is not critical, since inference is accurate even with slightly noisy data.

**Figure 5.**
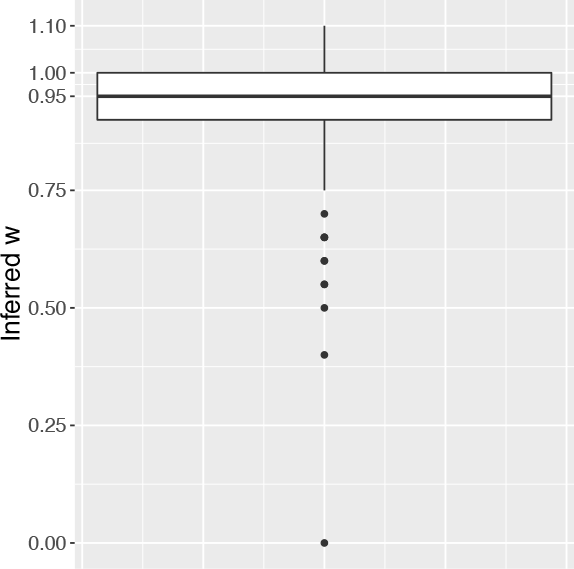
Range of fitness values inferred at the end of the iteration.

## 5 Introducing noise into the analysis

In many setups, population parameters may be imprecisely estimated. For example, microbial population size may actually be smaller or larger by an order of magnitude due to either a simple experimental error or due to inherent difficulty to infer it. Therefore, it is of great interest to see how FITS is affected by incorrect input parameter values.

We took a subset of the simulated datasets used for demonstrating the accuracy of FITS, and ran the fitness inference again, intentionally using wrong values for the population size (Figure 6A) and mutation rate (Figure 6B). When focusing on incorrect population size used as input, we observed that our inference of fitness remained quite robust for both neutral and advantageous alleles. Fitness was slightly underestimated in these cases when the input population size was an order of magnitude lower than the true value, making FITS more conservative in estimation of adaptive alleles. For lethal alleles, we observed very accurate inference even when the input population size was too high. However, FITS overestimated the fitness of lethal alleles when the input population size was too low: lethal alleles were on average estimated as having a fitness of 0.27±0.11 when the input population size was half of the real size, and 0.83±0.051 when the input population size was one tenth of the real size.

**Figure 6.**
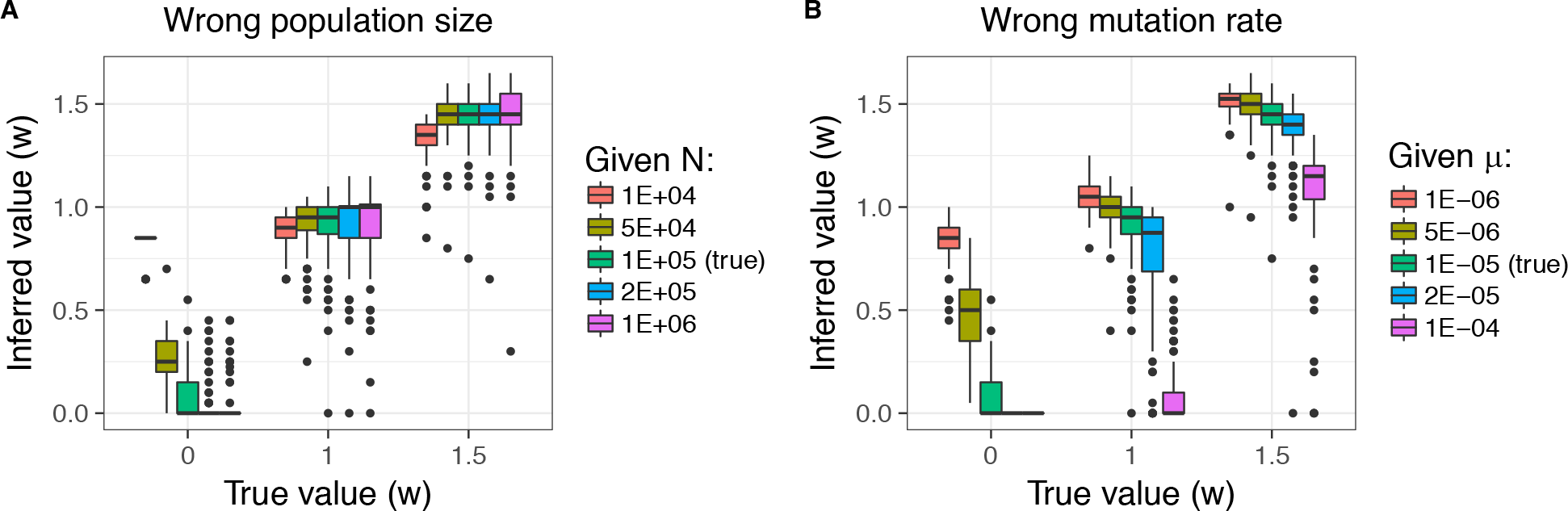
Effect of incorrect values inputted to FITS on accuracy of fitness inference. Boxplots are as described in Fig. 3. (A) Incorrect input population size, while mutation rate and fitness are fixed to true values. (B) Incorrect input mutation rate, while population size and fitness are fixed to true values. See main text for details.

On the other hand, inputting wrong mutation rates to FITS had a more complex effect on the accuracy. For neutral and advantageous alleles, giving as input an extremely low mutation rate had little effect, and FITS quite accurately inferred the fitness values in these cases. However, too high an input for the mutation rates caused FITS to strongly underestimate the fitness values of neutral and advantageous alleles. In fact, neutral alleles were often estimated as lethal if the mutation rate given as input was an order of magnitude higher than the real value (0.083±0.16). This is consistent with predictions from mutation-selection balance theory: a lethal allele is expected to be maintained at a frequency of the mutation rate. If the erroneously given mutation rate is very high, neutral alleles will remain below this mutation rate over the short time frame simulated, and will hence be classified as lethal. While this is a critical point to notice, it still emphasizes that FITS remains conservative for advantageous alleles and will not report false positives. Similar to the case with inaccurate population sizes, fitness of lethal alleles tends to also be overestimated when too low a mutation rate is given as input. The dangers of inference with incorrect input values are summarized generally in Table 3.

**Table 3.**
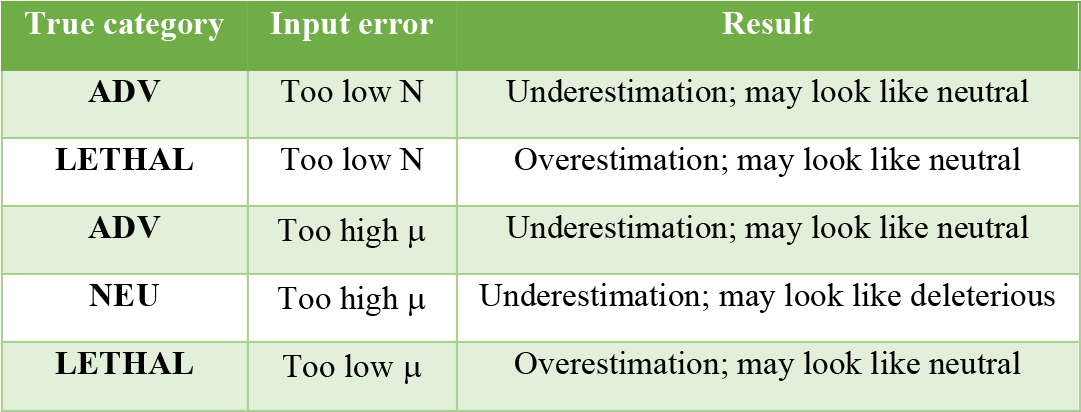
Summary of possible inference errors obtained when inputting incorrect mutation rates or population sizes.

In summary, when the mutation rate or population size are twice as high or twice as low as the real value (i.e., same order of magnitude), the inference of FITS is still quite robust. However, when FITS receives as input a parameter that is an order of magnitude higher or lower than the real value, this has a pronounced effect on inference of lethal (and presumably also non-lethal deleterious) alleles. Importantly, FITS remains conservative with advantageous alleles and tends to not overestimate their fitness.

## 6 Case Study – OPV2 quadrallelic analysis

We next set out to use FITS to analyze empirical data obtained from NGS of oral poliovirus type 2 (OPV2) that we have previously performed (Stern et al., 2017). Briefly, OPV2 was serially passaged (Figure 1D) at 39.5°C for seven passages, corresponding to fourteen generations. During the experiment, a population of *N* = 10^6^ infectious virus particles (Plaque Forming Unit or PFU) were seeded onto about 10^7^ cells grown in tissue culture. Each passage was sequenced using highly accurate CirSeq sequencing (Acevedo et al., 2014), allowing the detection of mutations at a frequency as low as 10^−6^. Coverage (number of reads covering a locus) spanned between 10^5^ and 10^6^ across all sequenced passages. In the analysis described below, we applied a filtering step and removed allele frequencies inferred with low confidence (based on geometric probability), which are typically alleles with very low read counts (Stern et al., 2017).

We used FITS as follows: first, we ran FITS on each locus (independently) to infer the fitness of each allele. Here we assumed a conservative population size of 10^5^. Mutation rates given as input were based on estimates obtained previously based on linear regression of synonymous mutation frequencies, under the assumption that they are mostly neutral (Stern et al. 2017). Next, in order to test how FITS infers mutation rates, we ran FITS independently on each presumably neutral synonymous mutation. Accordingly, FITS was given *w* = 1 as input, and once again N was set to 10^5^. Results were compared to the linear regression results obtained previously. Finally, we used FITS to also infer the population size, by running FITS independently on each (once again presumably neutral) synonymous mutation. Accordingly, FITS was given *w* = 1 as input, and the mutation rates were set to the values obtained from the linear regression. The results of all these analyses are described below.

### 6.1 Inferring fitness of each mutation in the genome of OPV2

We first ran an analysis with FITS using the biallelic model, applied to transition mutations only. Next, we ran an analysis using the quadrallelic model, applied to loci where all four nucleotides were observed. FITS was run on each locus independently. The results of both analyses give the distribution of fitness effects (DFE; Figure 7A) of the virus. Notably, this is a unique in-depth view of genomic evolution, enabled due to (a) the very high sequencing depth in the experiment, (b) the very high rate of mutation of the viral populations, and (c) the highly accurate sequencing approach used. Our results show a clear difference in the distribution of fitness effects obtained with transitions versus transitions+transversions. Transversions tended to be enriched with more non-lethal deleterious variants, whereas transitions were far less deleterious in general (Figure 7A). Indeed, this is in line with the genetic code structure, since transitions will more often create synonymous mutations, and when creating non-synonymous mutations, transitions often create more similar amino-acids (Sella and Ardell, 2002).

**Figure 7.**
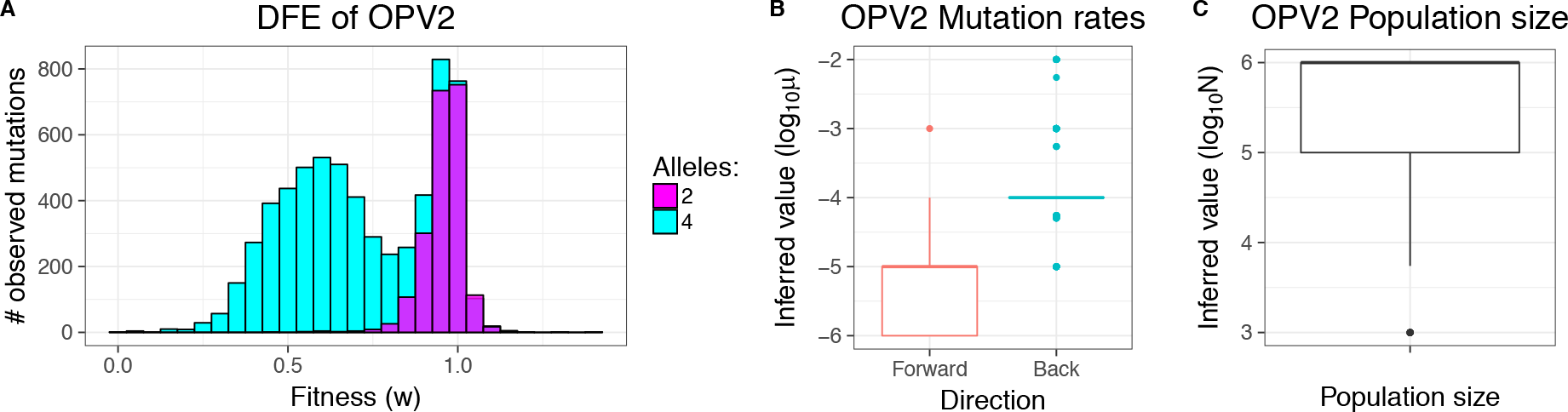
Inferred parameters for OPV2: (A) The distribution of fitness effects (DFE) across all loci in the genome of OPV2 based on FITS under a biallelic or quadrallelic model. Note that using the quadrallelic model (transitions + transversions) extends the one of the biallelic model (transition mutations only), revealing additional deleterious alleles. Low confidence allele frequencies were filtered out, leading to the observed depletion in lethal and highly deleterious alleles. (B) Inferred transition mutation rates for OPV2 across all synonymous loci. (C) Inferred population size for OPV2 in the described experiment, across all synonymous loci.

### 6.2 Inferring the population-wide mutation rates and population size of OPV2

We next set out to infer the mutation rate for the transition mutations of OPV2. Only loci where a synonymous transition mutation was observed were used for the analysis. FITS inferred both the back and forward mutation rates independently at each locus, and the values are presented as a boxplot in Figure 5B. The forward mutation rate estimates ranged mostly between 10^−6^ and 10^−5^, with a median value of 10^−5^. This is in agreement with the transition mutation rates we inferred previously using linear regression (Stern et al., 2017), which spanned 5 × 10^−6^ − 1 × 10^−5^. In a similar manner, we inferred the population size of the virus, based on independent inference of the population size at each locus where a synonymous transition mutation was observed. Inferred population sizes ranged mostly between 10^5^ and 10^6^ (Figure 7C), which is in agreement with the experimental protocol used to seed 10^e^ plaque forming units (PFUs) at each passage (Stern et al., 2017).

## 7 Discussion

We have developed FITS, a generic method that allows analyzing time-series data, and inferring the key parameters that shaped the evolutionary trajectory of an allele in an experiment or in real life settings. The program was designed with our recent evolutionary experiments of RNA virus populations in mind (Stern et al., 2017). These experiments monitor a population of viruses that begins as a clonal entity, and accumulates genetic diversity rapidly due to the high rate of mutation of the RNA viruses. In the initial setup of the experiment, genetic drift plays a prominent role, since mutations are born and present at low copy numbers. However, FITS is generic enough to be used to analyze other types of data, essentially any evolutionary experiment that tracks the population frequency of a trait over time.

Some of the key advantages of FITS include the fact that through the simulations, FITS is able to mimic the true biology of an allele as it is born and spreads in the population. FITS incorporates the mutation rate, allowing the introduction of new mutations along time. This is especially vital for new arising mutations. Moreover, by directly modeling stochastic effects, FITS takes into account fluctuations in allele frequencies, which may be quite prominent in the first few generations. Reassuringly, our results show a very high level of accuracy in simulated data. In fact, even when the parameters are erroneous to some extent, FITS remains robust under some conditions. We hope the fact that we have delineated the conditions where FITS tends to err, will allow users to be cautious when interpreting the result.

It is important to delineate the limitations of FITS, which represent the assumptions of the Wright-Fisher model used for simulations and the entire framework used. First, FITS assumes that loci evolve independently, and hence each locus is analyzed separately; accordingly, phenomenon such as linkage, or epitasis, are not taken into account. This is a relatively realistic assumption when considering evolutionary experiments spanning a short time frame, since typically only one mutation per genome will be present. However even so, and particularly for longer time-spans, this assumption is likely to be violated (Martin and Roques, 2016). We note that alleviating this assumption by considering all possible interactions among sites would be extremely computationally burdensome, and future work will be required to address non-independent evolution among sites. A second limitation of FITS has to do with the amount of information present in the experiment. Our simulation results showed that when the copy number of the allele is low, as reflected by a low *Nμ*, there is not enough information to infer fitness with FITS. Furthermore, when the data is very limited, for example when only one locus and two time-points are given for inference, FITS may also be unreliable (data not shown). Importantly, one of the features of FITS is the ability of the program to detect unreliable inference, both when *Nμ* is too low, and also when the posterior distribution yields no additional information over the prior distribution, and to output a warning to the user.

We note that FITS is designed to infer parameter values regarding one specific locus in a genome. However, FITS is clearly more robust when multiple loci are used to infer a specific parameter that is supposedly shared across many loci, such as the population size or mutation rate of a specific category of sites. This is also true for fitness – while naturally a user may be interested in the fitness of one particular allele, fitness inferred for a class of alleles (for example a particular type of non-synonymous mutations) will likely yield more robust results. Currently, FITS supports analyzing multiple loci by analyzing them independently and obtaining a distribution of inferred values, from which it is possible to obtain to obtain bounds on the desired parameter.

To summarize, FITS is a generic tool that may be used for inferring mutational fitness, mutation rates, or population size. To the best of our knowledge this is the first available tool for inferring mutation rates and population sizes (but see (Ferrer-Admetlla et al., 2016) who allow inference of population-size scaled selection), and the first highly user-friendly tool for inferring fitness of virus population. Finally, we have made a great effort in making FITS highly user friendly and intuitive for understanding, hopefully making this another milestone in making the tools of contemporary computational biology available to all virologists.

## 8 Acknowledgements

We thank Eli Levy Karin and Stern Lab group members for commenting on the manuscript. This work was supported in part by the Israeli Science Foundation (grant number 1333/16), and by a fellowship from the Edmond J. Safra Center for Bioinformatics at Tel-Aviv University.

